# INO-4800 DNA Vaccine Induces Neutralizing Antibodies and T cell Activity Against Global SARS-CoV-2 Variants

**DOI:** 10.1101/2021.04.14.439719

**Authors:** Viviane M. Andrade, Aaron Christensen-Quick, Joseph Agnes, Jared Tur, Charles Reed, Richa Kalia, Idania Marrero, Dustin Elwood, Katherine Schultheis, Mansi Purwar, Emma Reuschel, Trevor McMullan, Patrick Pezzoli, Kim Kraynyak, Albert Sylvester, Mammen P. Mammen, Pablo Tebas, J. Joseph Kim, David B. Weiner, Trevor R.F. Smith, Stephanie J. Ramos, Laurent M. Humeau, Jean D. Boyer, Kate E. Broderick

**Affiliations:** Inovio Pharmaceuticals, Plymouth Meeting, PA, 19462, USA; Vaccine and Immunotherapy Center, Wistar Institute, Philadelphia, PA 19104, USA; Hospital of the University of Pennsylvania, Philadelphia, PA, USA

## Abstract

Global surveillance has identified emerging SARS-CoV-2 variants of concern (VOC) associated with broadened host specificity, pathogenicity, and immune evasion to vaccine induced immunity. Here we compared humoral and cellular responses against SARS-CoV-2 VOC in subjects immunized with the DNA vaccine, INO-4800. INO-4800 vaccination induced neutralizing antibodies against all variants tested, with reduced levels detected against B.1.351. IFNγ T cell responses were fully maintained against all variants tested.

## Main Text

SARS-CoV-2, the causative agent of the COVID-19 pandemic, continues to cause unprecedented levels of mortality and socioeconomic burden. Concerningly, virus surveillance shows the global spread of novel SARS-CoV-2 variants, which are more infectious and display increased transmissibility and pathology [1-3]. Some of these variants of concern (VOC) contain mutations in the Spike protein receptor binding domain (RBD), the region which interacts with the host ACE2 receptor, and to which many SARS-CoV-2 neutralizing antibodies target. The B.1.1.7 lineage, the first emerging VOC reported in the United Kingdom, contains the key N501Y and D614G mutations and the 69-70 deletion in the RBD and/or S1 regions and exhibits increased transmissibility which is associated with increased pathogenesis, but does not appear to significantly evade neutralizing antibody responses generated by current vaccines approved for use [4, 5]. The B.1.351 (first reported in South Africa) and P.1 (first reported in Brazil) lineages have additional mutations, including E484K in the RBD region [6-8], as well as unique changes in position K417. Notably, sera isolated from convalescent individuals and vaccinees exposed to the wild-type (WT) Spike protein sequence have shown significantly lower levels of neutralizing activity against the B.1.351 and P.1 variants [8-12].

INO-4800 is a SARS-CoV-2 Spike DNA-based vaccine that is delivered intradermally followed by electroporation (EP) using CELLECTRA^®^ 2000 and is currently undergoing clinical development. In a Phase 1 clinical trial, INO-4800 vaccination induced a balanced immune response characterized by both functional antibody and T cell responses in vaccinated subjects [13]. Both humoral and cellular immune responses have been shown to be important components of protection against betacoronaviruses [14-16]. In the present study, we have assessed the humoral and T cell responses against SARS-CoV-2 B.1.1.7, B.1.351 and P.1 variants elicited after INO-4800 vaccination (Figure S1A).

In INO-4800 vaccinated subjects, serum IgG antibody binding titers to SARS-CoV-2 full-length Spike proteins were evaluated by ELISA using proteins specific for B.1.1.7, B.1.351, and P.1 variants (Fig. 1A, and S1A). IgG binding titers were not negatively impacted between WT and B.1.1.7 or B.1.351 variants. An average 1.9-fold reduction was observed for the P.1 variant in subjects tested at week 8 after receiving two doses of INO-4800 (Fig. 1A).

**Figure 1.**
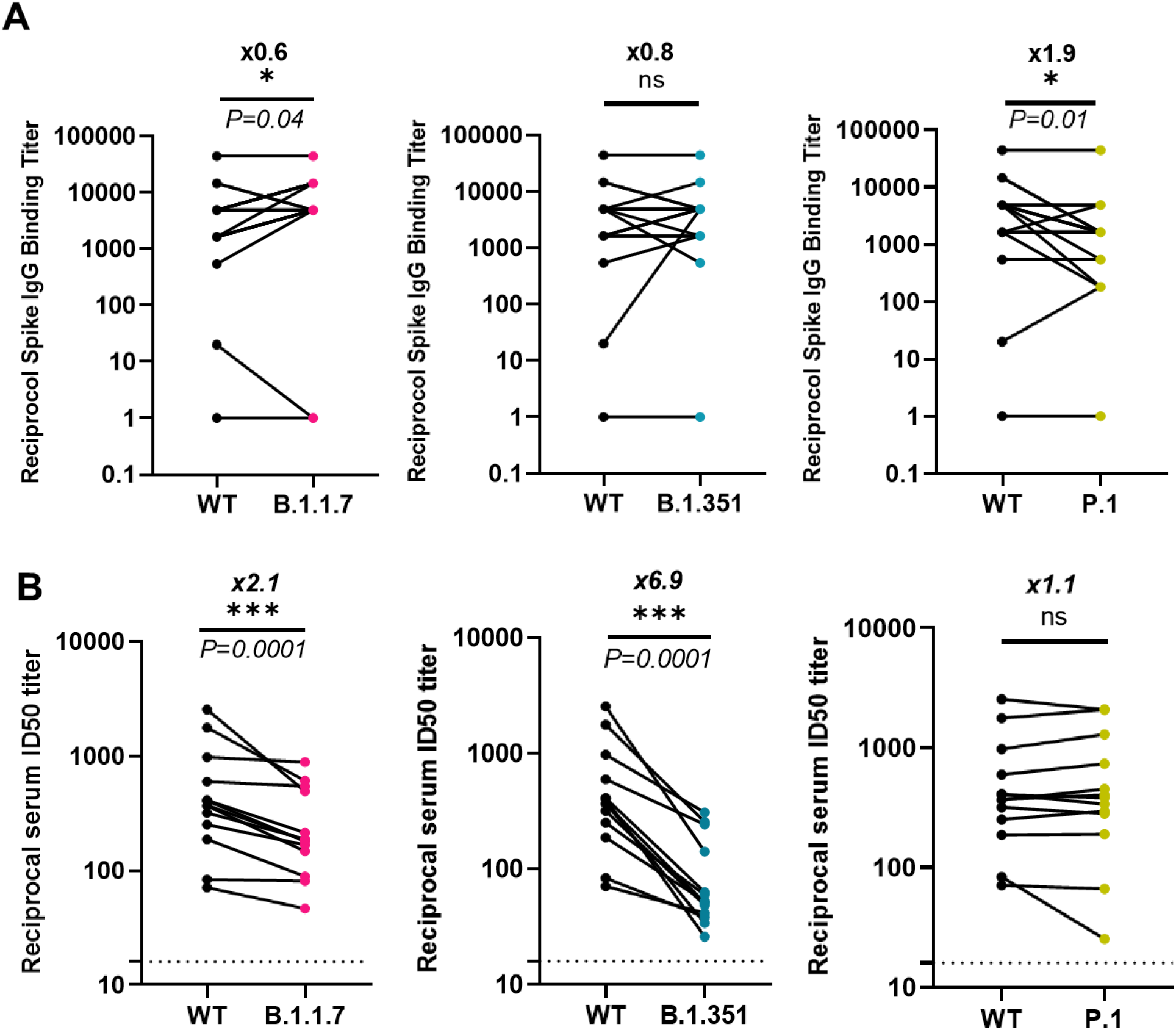
Humoral antibody cross-reactivity responses against SARS-CoV-2 variants. **a)** Sera from Phase 1 INO-4800 vaccinees were assessed by ELISA for IgG binding to WT, B.1.1.7, B.1.351, and P.1 variant Spike proteins (S1 and S2). Data points indicate endpoint titers for an individual study sample *(n=4, 0*.*5 mg vaccine dose; n = 5, 1*.*0 mg; n = 11, 2*.*0 mg)* and were calculated as the titer that exhibited an OD 3 SD above baseline. **b)** SARS-CoV-2 pseudovirus neutralization ID50 titers for sera samples from 13 Phase 1 INO-4800 vaccinees comparing WT against B.1.1.7, B.1.351 and P.1. Each data point represents the mean of technical duplicates for each individual (n=1, 0.5 mg vaccine dose; n = 4, 1.0 mg; n = 8, 2.0 mg). Dotted lines indicate the limit of detection of 16. ns – not significant, *P < 0.05, ***P < 0.0001 (Wilcoxon signed-rank test).

We performed a SARS-CoV-2 pseudovirus neutralization assay using sera collected from thirteen subjects two weeks after administration of a third dose of either 0.5 mg, 1 mg, or 2 mg of INO-4800 (Table S1). Neutralizing activity was detected against WT and the emerging variants in all serum samples tested (Fig. 1B). The mean ID50 titers for the WT, B.1.1.7, B.1.351 and P.1. were 643, 295, 105, and 664, respectively (Table S1). Compared to WT, there was a 2.1 and 6.9-fold reduction for B.1.1.7 and B.1.351, respectively, while there was no difference between WT and the P.1 variant. These results are consistent with other recent studies, which have demonstrated a significant reduction in neutralizing activity in vaccinated individuals towards the B.1.351 (≥6-fold reduction), while the B.1.1.7 lineage has demonstrated a reduced activity of 2-fold or less [4, 7-9, 17]. Strikingly, while the P.1 strain presents with similar RBD mutations as B.1.351, we did not observe any reduction in neutralizing activity compared to the WT strain in INO-4800 vaccinated individuals [18, 19]. The P.1 lineage has similar changes in the RBD to the B.1.351 lineage as they contain the N501Y mutation found in the B.1.1.7 lineage and identical E484K mutations. However, they have similar but different mutations at position K417, with a change to T for the P.1 and to N for the B.1.351 lineage. While both are changes to polar uncharged side chains, the B1.351 N mutation carries an additional amide group. It is possible that this position and this change allows for the impact of neutralizing responses observed.

Recent reports have supported an important role for T cell immunity in protecting against COVID-19 in the absence of antibodies to SARS-CoV-2 [20, 21]. Through optimized DNA construct design, combined with intradermal enhanced delivery, Inovio’s DNA platform technology has been demonstrated to drive balanced humoral and cellular immune responses to a wide range of infectious disease and tumor antigen targets [22-24]. We therefore compared cellular immune responses to WT and SARS-CoV-2 Spike variants elicited by INO-4800 vaccination. Peripheral blood mononuclear cells (PBMCs) isolated from ten subjects at week 8 after receiving their second dose of INO-4800 were stimulated with WT, B.1.1.7, B.1.351, or P.1 Spike peptides and cellular responses were measured by IFNγ ELISpot assay. We observed strikingly similar cellular responses to WT (median = 82.2 IFNγ spot-forming units [SFUs]/10^6^ PBMCs, IQR = 58.9-205.3), B.1.1.7 (median = 79.4, IQR = 38.9-179.7), B.1.351 (median = 80.0, IQR = 40.0-208.6) and P.1 (median = 78.3, IQR = 53.1-177.8) Spike peptides (Fig. 2). This is consistent with published results showing that, compared with neutralizing antibody responses, cellular immunity is relatively unimpaired by the current variants of concern [25]. Here, we show that T cell responses are consistently maintained between WT and all SARS-CoV-2 variants tested, including B.1.351 and P.1. Cells stimulated with peptides against these variants generated IFNγ responses as well as cytokines associated with CD8+ cytotoxic T cell responses (data not shown).

**Figure 2.**
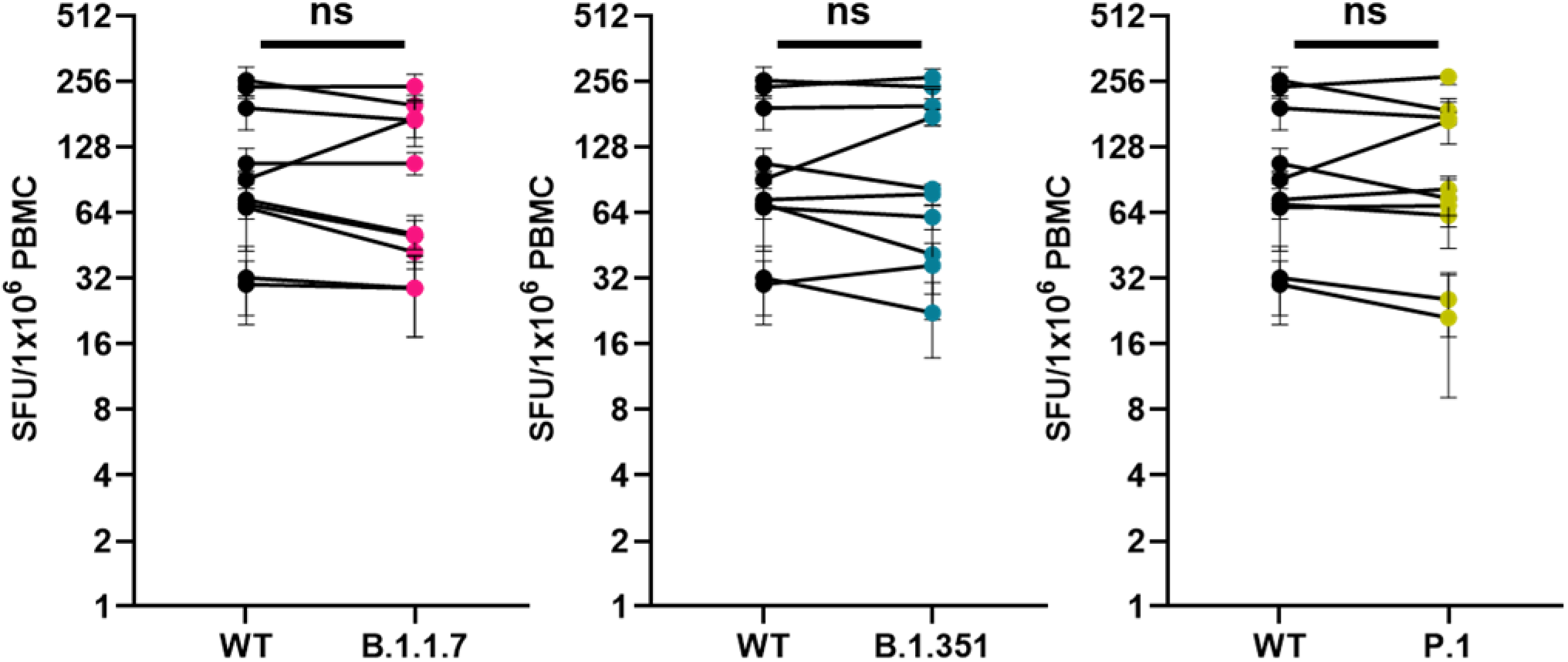
INO-4800 cellular immune response against SARS-CoV-2 variants. PBMCs from 10 Phase 1 subjects were collected 8 weeks after receiving the second dose of INO-4800 *(n = 5, 1*.*0 mg; n = 5, 2*.*0 mg*). PBMCs were treated with peptide pools spanning the entire Spike proteins of the WT, B.1.1.7, B.1.351, or P.1 variants and cellular responses were measured by IFNγ ELISpot assay. Mean +/- SEM IFNγ SFUs/million PBMCs of experimental triplicates are shown. ns - not significant (Wilcoxon signed-rank test).

There is a growing concern regarding the protective efficacy of vaccines recently approved for emergency use and those vaccines currently in development against the variants of concern. Recent studies have shown a reduction in serum neutralization levels against B.1.351 and P.1 [4, 7, 18], with minimal impact against B.1.1.7 [17]. In addition, vaccine trials have shown a considerable reduction in protective efficacy against B.1.351 [11, 12, 26]. Interestingly, a small reduction in protective efficacy is observed in Latin American countries including Brazil, suggesting that P.1 may have emerged around the time of clinical trials were in effect [26]. Here we report the neutralizing antibody and T cell activity measured in INO-4800 vaccinated subjects against emerging SARS-CoV-2 variants first detected in the United Kingdom, South Africa, and Brazil. The neutralization levels against B.1.351 and B.1.1.7 for the INO-4800 SARS-CoV-2 Spike DNA vaccine are consistent with previous reports of subjects receiving vaccines encoding for the ancestral Spike protein [7, 27]. Surprisingly, despite recent reports showing a reduction in neutralizing activity against the P.1 variant [8, 18], INO-4800 generates robust neutralizing antibodies at levels comparable to the WT strain. INO-4800 induces cross-reactive T cell responses against B.1.1.7, B.1.351, and P.1 variants that are comparable to the WT strain. Taken together, these data demonstrate maintenance of one or both cellular and humoral arms of the immune response against emerging SARS-CoV-2 variants for the INO-4800 vaccine, which will likely be critical factors to impact the ongoing COVID-19 pandemic.

## Methods

### Clinical Trial Subject Samples

Serum and PBMC samples were acquired from participants of the phase I INO-4800 clinical trial (NCT04336410) described previously [13]. The trial has since been expanded to include participants of 51-64 and 64+ years of age as separate groups in addition to the original 18-50 age group. A 0.5 mg dose group was also added. Sera from 20 subjects out of the 120 total study participants were selected for analysis on variant Spike protein binding ELISAs and variant pseudovirus neutralization assays. The samples analyzed by pseudovirus neutralization assay were collected from subjects two weeks after a third dose of INO-4800, and the samples used for other ELISA and ELISpot were collected after two doses.

### Antigen Binding ELISA

Binding ELISAs were performed as described previously[13], except different variants of SARS-CoV-2 S1+S2 proteins were used for plate coating. The S1+S2 wild-type Spike protein (Acro Biosystems #SPN-C52H8) contained amino acids 16-1213 of the full Spike protein (Accession #QHD43416.1) with R683A and R685A mutations to eliminate the furin cleavage site. The B.1.1.7, B.1.351, and P.1 S1+S2 variant proteins (Acro Biosystems #SPN-C52Hc,#SPN-C52H6, and #SPN-C52Hg, respectively) additionally contained the following proline substitutions for trimeric protein stabilization: F817P, A892P, A899P, A942P, K986P, and V987P. The B.1.1.7 protein contained the following variant-specific amino acid substitutions: HV69-70del, Y144del, N501Y, A570D, D614G, P681H, T716I, S982A, D1118H; the B.1.351 protein contained the following substitutions: L18F, D80A, D215G, R246I, K417N, E484K, N501Y, D614G, A701V; and the P.1 protein contained the following: L18F,T20N,P26S, D138Y,R190S,K417T,E484K,N501Y,D614G,H655Y,T1027I,V1176F. Assay plates were coated using 100 µL of 2 µg/mL of protein.

### SARS-CoV-2 Pseudovirus Production

SARS-CoV-2 pseudovirus stocks encoding for the WT, B.1.1.7, B.1.351 or P.1 Spike protein were produced using HEK 293T cells transfected with Lipofectamine 3000 (ThermoFisher) using IgE-SARS-CoV-2 S plasmid variants (Genscript) co-transfected with pNL4-3.Luc.R-E-plasmid (NIH AIDS reagent) at a 1:8 ratio. 72h post transfection, supernatants were collected, steri-filtered (Millipore Sigma), and aliquoted for storage at -80°C.

### SARS-CoV-2 Pseudoviral Neutralization Assay

CHO cells stably expressing ACE2 (ACE2-CHOs) were used as target cells plated at 10,000 cells/well. SARS-CoV-2 pseudovirus were tittered to yield greater than 30 times the cells only control relative luminescence units (RLU) after 72h of infection. Sera from 13 INO-4800 vaccinated subjects were heat inactivated and serially diluted two-fold starting at 1:16 dilution. Sera were incubated with SARS-CoV-2 pseudovirus for 90 min at room temperature. After incubation, sera-pseudovirus mixture was added to ACE2-CHOs and allowed to incubate in a standard incubator (37% humidity, 5% CO2) for 72h. After 72h, cells were lysed using Bright-Glo(tm) Luciferase Assay (Promega) and RLU was measured using an automated luminometer. Neutralization titers (ID50) were calculated using GraphPad Prism 8 and defined as the reciprocal serum dilution at which RLU were reduced by 50% compared to RLU in virus control wells after subtraction of background RLU in cell control wells.

### SARS-CoV-2 Spike ELISpot assay

Peripheral mononuclear cells (PBMCs) were stimulated in vitro with 15-mer peptides (overlapping by 11 amino acids) spanning the full-length Spike protein sequence of the indicated variants. Variant peptide pools (JPT Pepmix^™^) included the following changes to match published deletions/mutation in each variant: B.1.1.7 variant (delta69-70, delta144, N501Y, A570D, D614G, P681H, T716I, S982A, D1118H), B.1.351 variant (L18F, D80A, D215G, delta242-244, R246I, K417N, E484K, N501Y, D614G, A701V); P.1 variant L18F, T20N, P26S, D138Y, R190S, K417T, E484K, N501Y, D614G, H655Y, T1027I, V1176F). Cells were incubated overnight with peptide pools at a concentration of 1 μg per ml per peptide in a precoated ELISpot plate, (MabTech, Human IFNγ ELISpot Plus). Cells were then washed off, and the plates were developed via a biotinylated anti-IFN-γ detection antibody followed by a streptavidin-enzyme conjugate resulting in visible spots. After plates were developed, spots were scanned and quantified using the CTL S6 Micro Analyzer (CTL) with *ImmunoCapture* and *ImmunoSpot* software. Values are shown as the background-subtracted average of measured triplicates. The ELISpot assay qualification determined that 12 spot forming units was the lower limit of detection. Thus, anything above this cutoff signal is an antigen specific cellular response.

### Statistical Methods

GraphPad Prism 8.1.2 (GraphPad Software, San Diego, USA) was used for graphical and statistical analysis of data sets. *P* values of <0.05 were considered statistically significant. A nonparametric two-tailed student t-test Wilcoxon signed-rank test was used to assess statistical significance in Figures 1 and 2.

## Data availability

The data that support the findings of this study are available from the corresponding authors upon reasonable request.

## ACKNOWLEDGEMENTS

We acknowledge the members of the Inovio Pharmaceuticals R&D department for significant technical assistance. This work is funded by Coalition for Epidemic Preparedness Innovations (CEPI).

## AUTHOR CONTRIBUTIONS

Conceptualization, V.M.A., S.J.R., J.D.B., K.E.B., T.R.F.S., J.J.K., L.M.H.; Methodology V.M.A., A.CQ., I.M., J.A., C.R., D.E., K.S., T.M.; Investigation; V.M.A., A.CQ., I.M., J.A., C.R., D.E., K.S., T.M. P.P.; Resources, M.P., E.R., P.T., P.P., D.B.W.; Writing-original draft preparation V.M.A., J.T., A.CQ., I.M., J.A., C.R., D.E., K.S., T.M.; Writing-review and editing, V.M.A., J.T., A.CQ., I.M., J.A., C.R., D.E., K.S., T.M; Project administration, S.J.R., J.D.B., K.E.B., T.R.F.S., J.J.K., L.M.H.

## ADDITIONAL INFORMATION

Supplementary information is available for this paper at (Link)

## Correspondence

and requests for materials should be addressed to Kate E. Broderick.

