## Supplemental Material for "INO-4800 DNA Vaccine Induces Neutralizing Antibodies and T cell Activity Against Global SARS-CoV-2 Variants"

### Slide 1
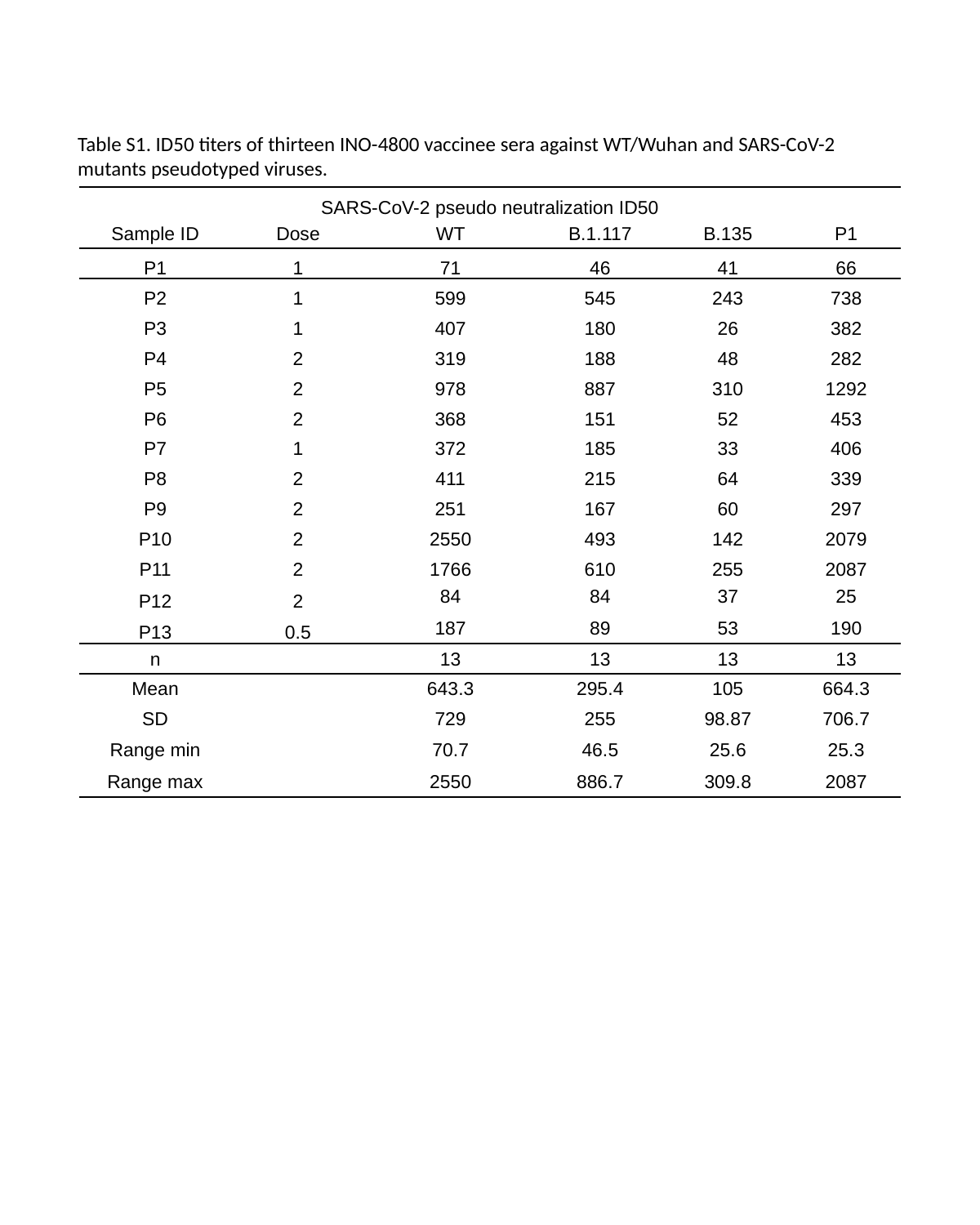

Table S1. ID50 titers of thirteen INO-4800 vaccinee sera against WT/Wuhan and SARS-CoV-2 mutants pseudotyped viruses.
| SARS-CoV-2 pseudo neutralization ID50 | | | | | |
| --- | --- | --- | --- | --- | --- |
| Sample ID | Dose | WT | B.1.117 | B.135 | P1 |
| P1 | 1 | 71 | 46 | 41 | 66 |
| P2 | 1 | 599 | 545 | 243 | 738 |
| P3 | 1 | 407 | 180 | 26 | 382 |
| P4 | 2 | 319 | 188 | 48 | 282 |
| P5 | 2 | 978 | 887 | 310 | 1292 |
| P6 | 2 | 368 | 151 | 52 | 453 |
| P7 | 1 | 372 | 185 | 33 | 406 |
| P8 | 2 | 411 | 215 | 64 | 339 |
| P9 | 2 | 251 | 167 | 60 | 297 |
| P10 | 2 | 2550 | 493 | 142 | 2079 |
| P11 | 2 | 1766 | 610 | 255 | 2087 |
| P12 | 2 | 84 | 84 | 37 | 25 |
| P13 | 0.5 | 187 | 89 | 53 | 190 |
| n | | 13 | 13 | 13 | 13 |
| Mean | | 643.3 | 295.4 | 105 | 664.3 |
| SD | | 729 | 255 | 98.87 | 706.7 |
| Range min | | 70.7 | 46.5 | 25.6 | 25.3 |
| Range max | | 2550 | 886.7 | 309.8 | 2087 |

### Slide 2
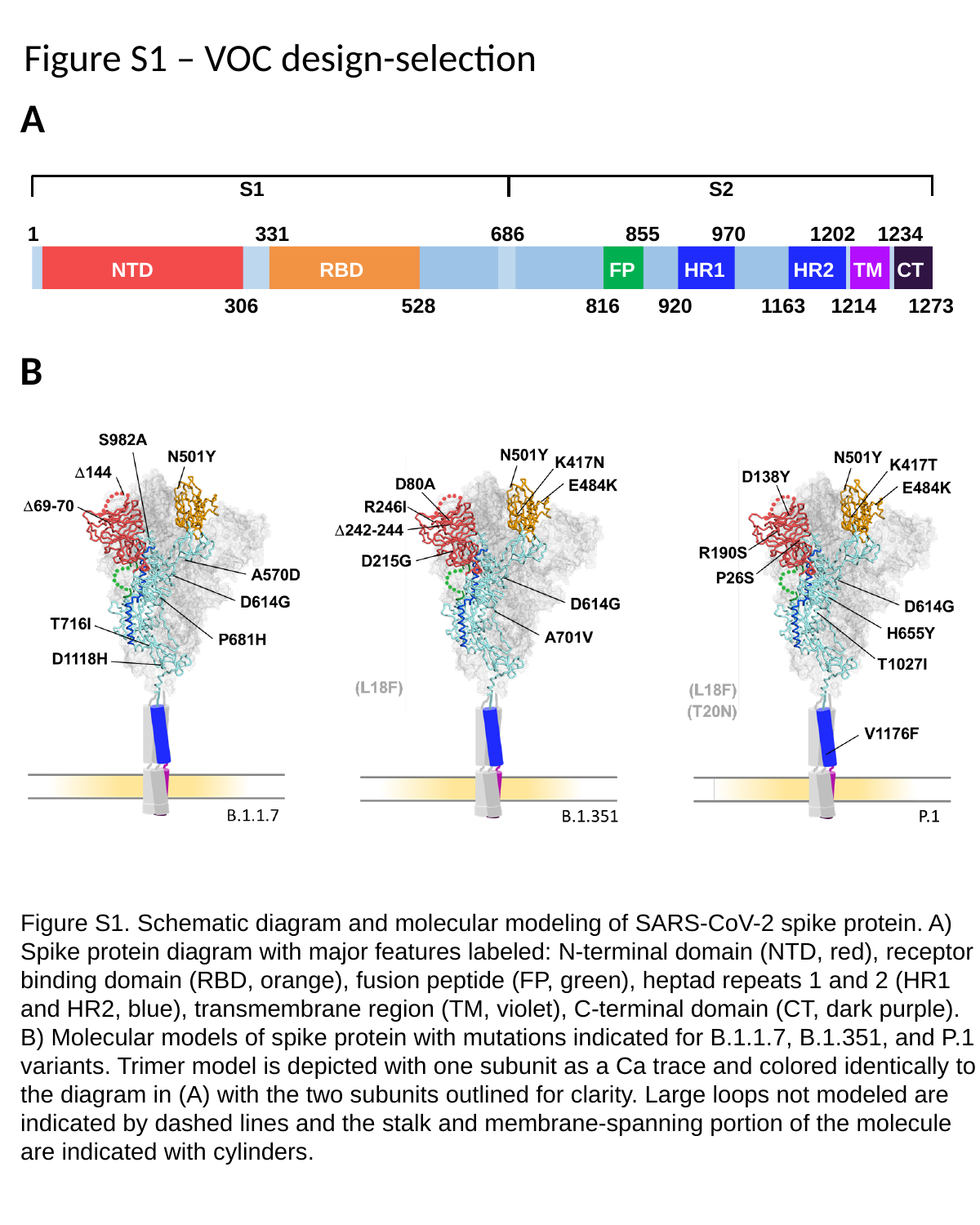

Figure S1 – VOC design-selection
A
S2
S1
1234
1
331
686
855
970
1202
NTD
RBD
FP
HR1
HR2
TM
CT
306
528
816
920
1163
1214
1273
B
Figure S1. Schematic diagram and molecular modeling of SARS-CoV-2 spike protein. A) Spike protein diagram with major features labeled: N-terminal domain (NTD, red), receptor binding domain (RBD, orange), fusion peptide (FP, green), heptad repeats 1 and 2 (HR1 and HR2, blue), transmembrane region (TM, violet), C-terminal domain (CT, dark purple). B) Molecular models of spike protein with mutations indicated for B.1.1.7, B.1.351, and P.1 variants. Trimer model is depicted with one subunit as a Ca trace and colored identically to the diagram in (A) with the two subunits outlined for clarity. Large loops not modeled are indicated by dashed lines and the stalk and membrane-spanning portion of the molecule are indicated with cylinders.
